# trio-sga: facilitating *de novo* assembly of highly heterozygous genomes with parent-child trios

**DOI:** 10.1101/051516

**Authors:** Milan Malinsky, Jared T. Simpson, Richard Durbin

**Affiliations:** Wellcome Trust Sanger Institute, Cambridge, CB10 1SA, UK.; Gurdon Institute and Department of Genetics, University of Cambridge, Cambridge, CB2 1QN, UK.; Ontario Institute for Cancer Research, Toronto, M5G 0A3, Canada.; Department of Computer Science, University of Toronto, Toronto, M5S 3G4 Canada.

## Abstract

**Motivation:** Most DNA sequence in diploid organisms is found in two copies, one contributed by the mother and the other by the father. The high density of differences between the maternally and paternally contributed sequences (heterozygous sites) in some organisms makes *de novo* genome assembly very challenging, even for algorithms specifically designed to deal with these cases. Therefore, various approaches, most commonly inbreeding in the laboratory, are used to reduce heterozygosity in genomic data prior to assembly. However, many species are not amenable to these techniques.

**Results:** We introduce `trio-sga`, a set of three algorithms designed to take advantage of mother-father-offspring trio sequencing to facilitate better quality genome assembly in organisms with moderate to high levels of heterozygosity. Two of the algorithms use haplotype phase information present in the trio data to eliminate the majority of heterozygous sites before the assembly commences. The third algorithm is designed to reduce sequencing costs by enabling the use of parents’ reads in the assembly of the genome of the offspring. We test these algorithms on a ‘simulated trio’ from four haploid datasets, and further demonstrate their performance by assembling three highly heterozygous *Heliconius* butterfly genomes. While the implementation of `trio-sga` is tuned towards Illumina-generated data, we note that the trio approach to reducing heterozygosity is likely to have cross-platform utility for *de novo* assembly.

## 1 Introduction

All existing genome sequencing technologies are limited by read lengths that are orders of magnitude shorter than whole chromosomes [see e.g. (Pendleton *et al.*, 2015)]. The goal of *de novo* genome assembly is to reconstruct the underlying genome sequence from the reads by finding and following overlaps between them. If error-free reads could be obtained from a single underlying genome sequence with randomly generated bases, the assembly task would be a trivial problem: a 50bp overlap between two reads would virtually guarantee that the two DNA sequences indeed originated from overlapping loci. In reality, however, almost all genome assembly projects are very challenging due to a combination of three problems: 1) DNA sequencing errors; 2) heterozygosity; 3) the highly repetitive nature of most eukaryotic genome sequences.

The relative importance of each problem is influenced by its severity. A high density of heterozygous sites makes genome assembly very challenging [see e.g. (Zhang *et al.*, 2012; Nystedt *et al.*, 2013)] and *de novo* assembly algorithms specifically designed for moderate-to-high levels of heterozygosity (Vinson *et al.*, 2005; Donmez and Brudno, 2011; Kajitani *et al.*, 2014) still cannot match the performance achieved in assemblies of homozygous strains, especially at the contig assembly level. Therefore, genome project teams often strive to reduce or even completely eliminate heterozygosity, either by using inbred laboratory strains [e.g. the *C. elegans* genome project (*C. elegans* Sequencing Consortium, 1998) and *D. melanogaster* genome project (Adams *et al.*, 2000)], or by obtaining a homozygous form of the organism by other laboratory techniques [e.g. the potato genome (Potato Genome Sequencing Consortium *et al.*, 2011)]. Using these approaches to reducing heterozygosity may be time consuming and sometimes not feasible, for example if the organism in question is resistant to inbreeding due to strong inbreeding depression (Charlesworth and Willis, 2009), or if the generation time is too long or the organism unsuitable for inbreeding in the laboratory. With ongoing projects to provide high quality genome assemblies for 10,000 vertebrate species (Genome 10K Community of Scientists, 2009), 5,000 arthropod species (i5K Consortium, 2013), and 7,000 (mainly marine) invertebrates (GIGA Community of Scientists *et al.*, 2014), there is a pressing need for new approaches to reducing heterozygosity for genome assembly. Genotyped individuals with familial relatedness and especially parents-child trios are commonly used for haplotype estimation (also known as ‘phasing’) in population genomics studies (B. L. Browning and S. R. Browning, 2009; O’Connell *et al.*, 2014). However, familial relatedness has to our knowledge never been used in the context of haplotype phasing during genome assembly, perhaps due to the computational complexity of this task. In this paper we take advantage of the phasing information available in one generation pedigrees comprising mother, father and one offspring. Such a trio of samples is often available or feasible to generate for species where long term inbreeding is not practical. We describe three efficient algorithms to facilitate haplotype phasing and assembly of highly heterozygous genomes. The algorithms, jointly referred to as `trio-sga`, are based around queries over an *FM-index*: a compressed index of all the reads (Ferragina and Manzini, 2000; Simpson and Durbin, 2010) and form an extension of the sga genome assembler (Simpson and Durbin, 2012). We demonstrate the performance of `trio-sga` by assembling a ‘simulated trio’ composed of reads from four haploid strains of *Schizosaccharomyces pombe*, and by assembling three highly heterozygous *Heliconius* butterfly genomes.

## 2 Methods

The way `trio-sga` algorithms fit into and extend the standard `sga` pipeline is depicted in Figure 1 (it also suggests how they could be used with or incorporated into other existing assembly software). The input into the `trio-sga` pipeline are three separate sets of DNA reads: reads from the mother, reads from the father, and reads from their offspring. During a typical workflow we build separate FM-indices for the three sets of reads. The aim is to assemble the reads from the offspring, taking advantage of haplotype phase information provided by data from the parents. If haplotype phase can be resolved at all heterozygous sites, the final products of our workflow are two separate assemblies of the offspring’s genome: an assembly of the haplotypes inherited from the mother (maternal) and an assembly of the haplotypes inherited from the father (paternal).

**Fig. 1:**
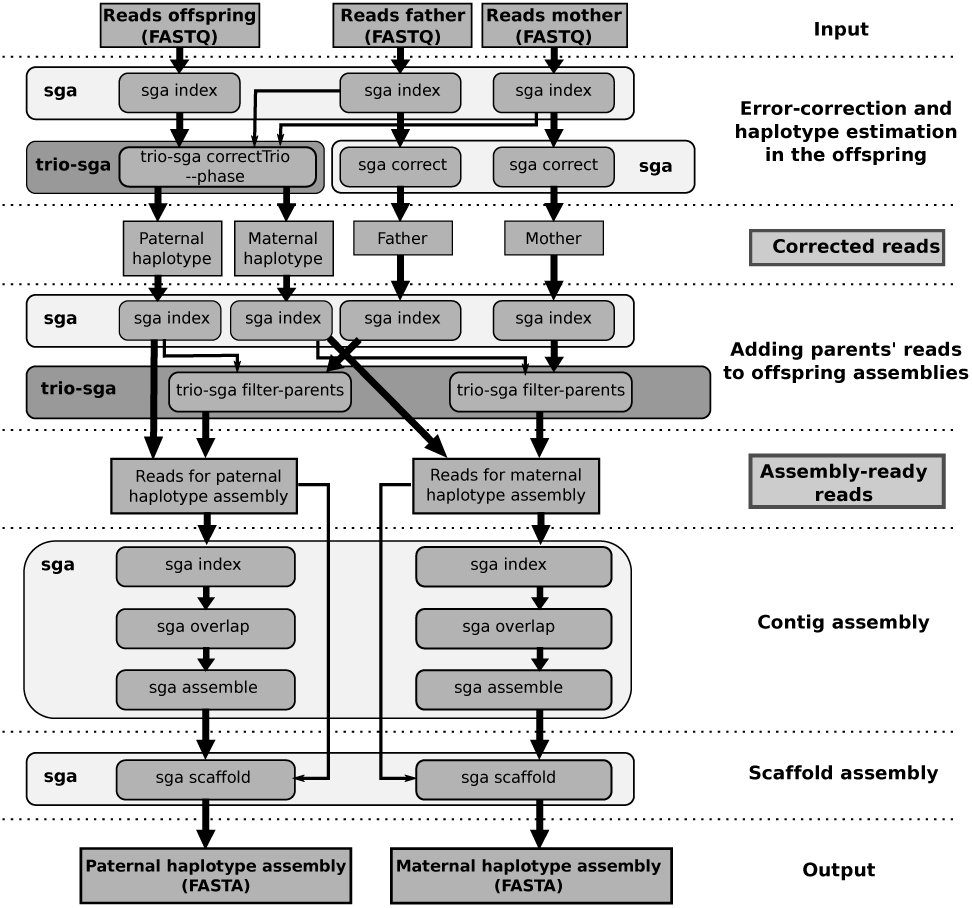
A diagram of a typical workflow showing how trio-sga algorithms fit into and extend the sga assembly pipeline. The main data flow is indicated by thick arrows. Thin arrows indicate inputs that are used to inform data processing algorithms but do not themselves form a part of the output.

The first `trio-sga` algorithm, implemented in `trio-sga cor-rectTrio`, uses trio information to improve the reliability of read error correction. Error correction used by sga and many other genome assemblers [see e.g. (R. R. Li *et al.*, 2010)] is based on *k*-mer frequencies-the number of occurrences of a sequences of length *k* in the overall dataset. This approach relies on the fact that the frequency distributions of correct and error-containing *k*-mers differ: the number of occurrences of error-containing *k*-mers is generally lower than the number of occurrences of *k*-mers that do not contain errors (Figure 2A,B). In practice, an occurrence threshold is set to distinguish between correct and error-containing *k*-mers. However, in most data sets (depending on the error rate) there is a ‘grey zone’ where the two distributions overlap (Figure 2B): low *k*-mer occurrences of correct sequence are possible due to low coverage and/or non-random sampling and high *k*-mer occurrences of error-containing *k*-mers, for example due to repeated (or systematic) errors. Using data from the parents helps to distinguish between error-containing and correct sequences within the grey zone and to prevent under-correcting (accepting as correct reads that contain errors) and over-correcting (‘fixing’ reads that are in fact correct). For example, if a *k*-mer fails the threshold in the offspring, but is present above threshold in one or both of the parents, it is unlikely to be an error and correction is not attempted.

**Fig. 2:**
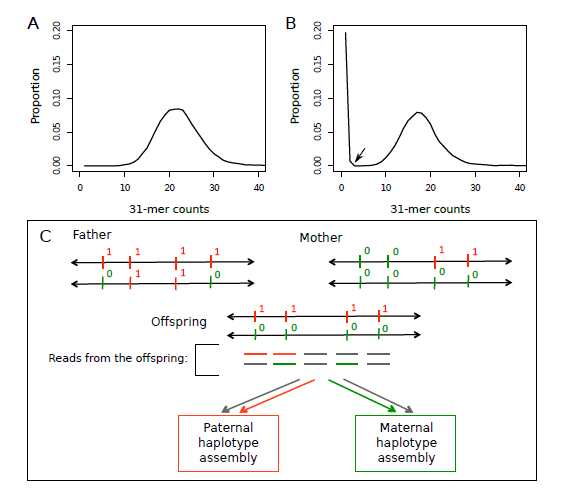
Trio aware error correction and read filtering/phasing. **(A)** The distribution of 31-mer counts in simulated 100bp error-free reads with 30X genome coverage of scaffold 1 of the *H. melpomene* assembly. **(B)** The same distribution is in (A) but reads were simulated with an Illumina HiSeq 2000 error profile, as implemented in the ART read simulator (Huang *et al.*, 2012). There are now many more 31-mers with low occurrences (<4); these are mainly errors, but there is a ‘grey zone’ (arrow), with 31-mers occurring 2-4 times being a mixture of correct and error-containing sequences. **(C)** An example region of the genome with four segregating sites. The offspring inherited a haplotype with four derived alleles (denoted as 1) from the father and a haplotype with four ancestral alleles from the mother. Reads (or read pairs) from the offspring that contain the first or the second segregating site and the derived allele can be phased and confidently assigned for paternal haplotype assembly (as indicated by red colour). Similarly, reads from DNA containing the second or the third segregating site and the ancestral allele can be phased and confidently assigned for maternal haplotype assembly (indicated by green colour). The remaining reads (grey) are assigned to both assemblies.

The second `trio-sga` algorithm, implemented in the—phase option of `trio-sga correctTrio`, filters the set of reads sequenced from the offspring in order to reduce heterozygosity. A conceptual overview of this algorithm is in Figure 2C. Reads are assumed to be error-free at this stage and to carry consistent phase information. Every *k*-mer of every read in the offspring is checked for phase-informative variants by counting the number of its occurrences in reads from each of the parents. Reads (or read-pairs) that carry *k*-mers present only in the mother are assigned to the maternal assembly and reads that carry *k*-mers present only in the father are assigned to the paternal assembly. Reads that carry *k*-mers present in both parents are assigned to both datasets.

The third algorithm has been designed to ‘fill’ regions of low coverage in the offspring by bringing in reads sequenced from the parents’ DNA, thus using the parents’ datasets to ‘assemble through’ these regions. In this way, the algorithm reduces the sequencing depth requirements and the costs of the genome assembly project. The algorithm works by checking every *k*-mer of every read in the father for consistency with the reads assigned for the offspring’s paternal haplotype assembly; all the consistent reads are then “brought in” to fill any coverage gaps. Reads from the mother are used in the same way for the maternal haplotype assembly.

The ‘simulated trio’ was obtained by combining publicly available paired-end 100bp Illumina reads from four strains of *S. pombe* (Jeffares *et al.*, 2015), as outlined in Figure 3. Briefly, each haploid yeast strain is treated as a parental haplotype: random subsets of reads from two strains were mixed together to represent each ‘parent’. The ‘offspring’ then inherited a separate random subset of reads from one strain from each ‘parent’ (recombination was not simulated, but would have negligible effect on `trio-sga` performance). The random sampling of reads was tuned to obtain 40X coverage per individual (i.e. 40X ‘father’, 40X ‘mother’, and 40X ‘offspring’). The specific read sets used are available from the European Nucleotide Archive (ENA) (Study: ERP000180; Accessions: ERS070898, ERS070920, ERS070926, ERS070966).

**Fig. 3:**
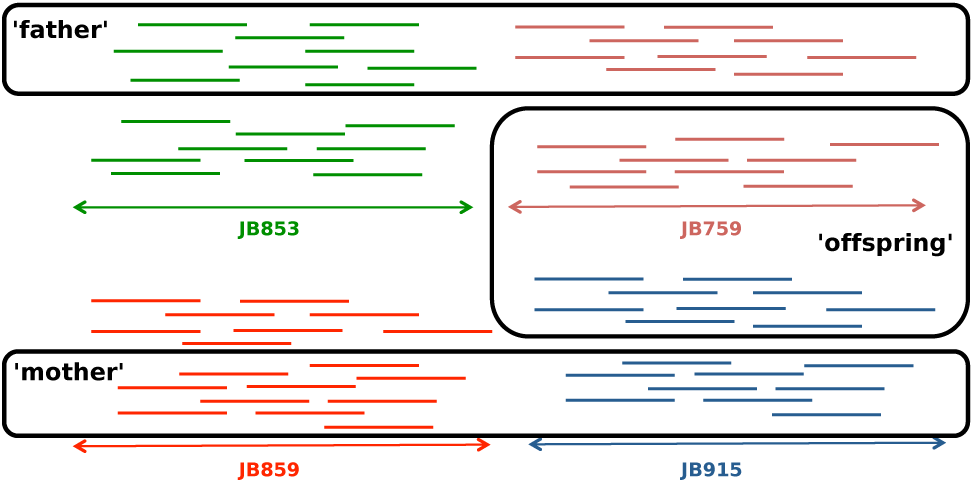
Yeast ‘simulated trio’ design. Reads from four haploid strains were used to represent the four ‘parental haplotypes’. The reads from strains JB853 and JB759 were combined to form the dataset from the ‘father’, and reads from strains JB859 and JB915 were combined to represent the ‘mother’. Non-overlapping random subsets of reads from strains JB759 and JB915 were then selected to form the ‘offspring’.

We generated new Illumina reads from parent-offspring trios of *Heliconius melpomene*, *Heliconius cydno*, and a laboratory cross between a *H. melpomene* father and *H. cydno* mother. Genomic libraries were prepared according to Illumina TruSeq HT protocol and sequenced using Illumina HiSeq v4 reagents to obtain 125bp paired-end reads with mean insert size of 300-500bp. All nine samples were multiplexed together and three lanes were sequenced, yielding ~124 billion base-pairs, corresponding to ~46X average genome coverage per individual (range ~40-50X). The raw read data are available from ENA (Study: ERP009507; Accessions: ERR926555-ERR926581).

The standard sga assemblies described in this manuscript were generated using the workflow from (Simpson and Durbin, 2012) and `trio-sga` assemblies using the workflow shown in Figure 1, with default parameters except the following: 1)`-k 41` was used for error correction in *Heliconius* assemblies; 2)`-r 10` was used for the sga assemble subprogram. The contig assemblies (output of sga assemble) were attempted with minimum overlap required between reads set to 65, 70, 75, and 85bp in yeast; and 70, 80, 90, 95, 100, 105, and 110bp in *Heliconius*. Then we choose the contig assembly with the highest N50 statistic. To obtain scaffolds we aligned the paired-end reads used for the trio assembly to the contigs (excluding contigs ≤200bp) using `bwa mem v0.7.10` (H. Li, 2013), and we required evidence from at least five pairs of reads before joining two contigs (`-n 5` parameter to the `sga-bam2de.pl` script). Finally, reads were re-mapped to the complete scaffold assembly (excluding scaffolds ≤500bp) and the proportion of properly-paired reads (a measure of assembly quality) obtained with `samtools flagstat v1.3` (H. Li, 2011).

## 3 Algorithms

The basic building block of all three algorithms is a simple query over an *FM-index*: counting the number of occurrences of a given *k*-mer and of its reverse complement in a read set. In what follows, *k*-mer occurrences in reads from the mother’s DNA are denoted C_M_(k), occurrences in reads from the father are denoted C_F_(k), and from the offspring C_O_(k).

### Algorithm 1: Trio-aware error correction: deciding whether to attempt correcting a *k*-mer

**Data**: FM-indices of mother, father, offspring reads: reads from the offspring

**Result**: Corrected offspring reads

1 // 

~~~
Initialise occurrence thresholds for *k*-mers in offspring, mother, and father FM-Indices
~~~

**Init:** set thresholds t_O_, t_M_, t_F_;

2 // 

~~~
Initialise indicator variables for ensuring haplotype phase consistency of corrected reads
~~~

**Init:** set **mc**=FALSE;set **fc**=FALSE;

3 **foreach** (read **R** *from the offspring)* **do**

4 **foreach** *(k-mer* **k** in **R**) **do**

5 if (C_F_(**k**) < *t_F_ and* C_M_(**k**) < t_M_) **then** increase t_O_

6 // 

~~~
Test the offspring threshold
~~~

7 **if** (C_O_(**k**) > t_O_) **then**

8 next; // 

~~~
Do not correct and move on to the next *k*-mer
~~~

9 else

10 // 

~~~
This *k*-mer failed offspring threshold - test it in the parents
~~~

11 **if** ((**mc** = = **fc**) *or* (**mc**= =TRUE *and* **fc**= = FALSE)) **then**

12 **if** (C_M_(**k**) > t_M_) **then** set **mc.temp**=TRUE;

13 **end**

14 **if** ((**mc**= = **fc**) or (**mc**= = FALSE *and* **fc**= =TRUE)) **then**

15 **if** *(C_F_(**k**) > t_F_)* **then** set **fc.temp**=TRUE;

16 **end if** (**mc.temp**= =TRUE *or* **fc.temp**= =TRUE) **then**

17 // 

~~~
Passed *k*-mer count threshold in the parental reads
~~~

18 set **mc** = **mc.temp**; set **fc** = **fc.temp:**

19 next;

20 // 

~~~
Do not correct and move on to the next *k*-mer
~~~

21 **end**

22 **end**

23 // 

~~~
Call the correction algorithm (not shown)
~~~

24 correction(**R.k**);

25 **end**

26 **end**

Line 5: if a *k*-mer found in the offspring does not occur (above threshold) in either parent, it is likely to be an error (or a de-novo mutation, but these are exceedingly rare compared with errors). Therefore, we increase the offspring *k*-mer occurrence threshold for this *k*-mer.

Algorithm 1 outlines how parents’ data are used in the decision on whether or not to attempt correction on a *k*-mer in the offspring. The correction algorithm itself is not shown as it is identical to the correction algorithm used by `sga` (Simpson and Durbin, 2012).

Algorithm 2 describes how parents’ data are used to reduce heterozygosity in the reads from the offspring by partitioning them into a maternal and paternal data sets.

### Algorithm 2: Filtering the set of reads sequenced from the offspring in order to reduce heterozygosity (assuming error-free reads)

**Data**: FM-indices of mother and father reads; reads from the offspring

**Result**: Two partially overlapping sets of offspring reads for paternal and maternal haplotype assembly.

1 **foreach**(*read* **R** *from the offspring*) **do**

2 set **inMother**=TRUE; set **inFather**=TRUE;

3 foreach (*k*-mer **k** in **R**) **do**

4 **if** (C_M_(k) = 0) **then**

5 **inMother**=FALSE;

6 **end**

7 **if** (C_F_(k) == 0) **then**

8 **inFather**=FALSE;

9 **end**

10 **end**

11 **if** (**inMother**==TRUE) **then** assign **R** to maternal assembly read set;

12 **if** (**inFather**==TRUE) **then** assign **R** to paternal assembly read set;

13 **end**

Algorithm 3 outlines how reads from the parents are checked for consistency with the reads assigned for the offspring’s paternal haplotype assembly.

### Algorithm 3: Check for consistency between error-corrected reads from the mother and error-corrected reads from the offspring’s maternal haplotype or reads from the father and the paternal haplotype.

**Data**: FM-indices of error-corrected reads from one parent and the corresponding haplotype in the offspring

**Result**: Reads from the parent that are consistent with the offspring read set and can be used to bridge coverage gaps

1 **foreach** (*read* **R** *from the parent*) **do**

2 **consistent**=TRUE;

3 **foreach** (*k-mer* **k** *in* **R**) **do**

4 **if** (C_*O*_(**k**) == 0) **then**

5 **consistent**=FALSE; break;

6 **end**

7 **end**

8 **if** (**consistent**) **then** mark **R** as consistent with the offspring;

9 **end**

## 4 Results and discussion

The performance of read filtering/phasing in reducing heterozygosity in the read sets is summarized in Figure 4. The estimated frequency of heterozygous sites in the ‘simulated yeast diploid’ offspring was 1 in 262 sites before and 1 in 569 sites after applying the `trio-sga` filtering (a reduction of 54%). In Heliconius butterfly trios, estimated heterozygosity decreased from 1 in 64 to 1 in 1371 sites (95.3% reduction) in *H. melpomene*; from 1 in 51 to 1 in 1098 sites in *H. cydno* (95.4% reduction); and from 1 in 34 to 1 in 1604 in the *H. cydno* x *H. melpomene* cross (97.9% reduction). The frequencies were estimated using the `sga-preqc` program (Simpson, 2014). Heterozygosity value for the haploid yeast strain (JB759) represents the misclassification rate (1 in 3043) observed in sga preqc estimates, resulting from mistaking error or repeat branches for heterozygous loci.

**Fig. 4:**
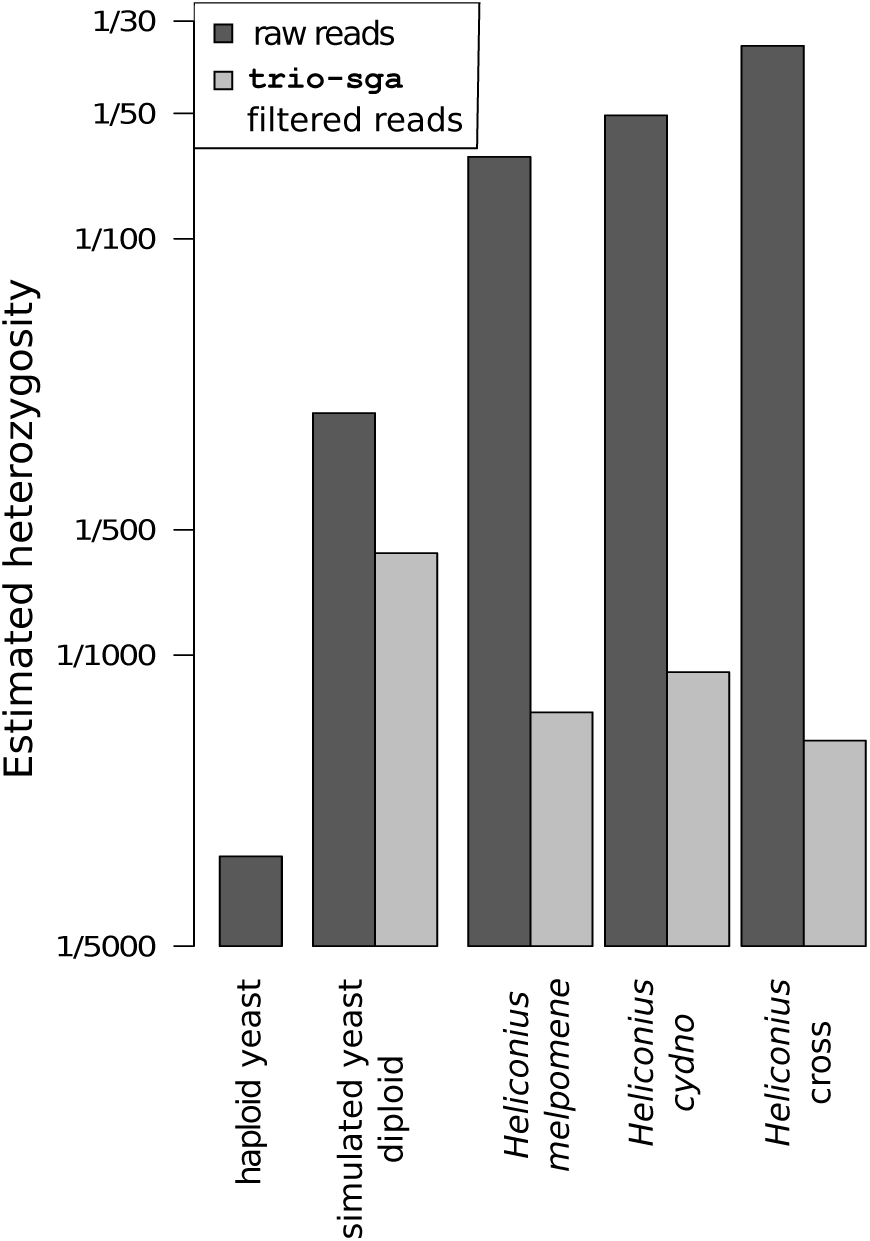
Read-filtering/phasing performance. The frequencies of heterozygous sites were estimated before `trio-sga` filtering in the offspring and after `trio-sga` filtering in the reads assigned for paternal haplotype assembly.

The results suggest that the higher the heterozygosity the better the `trio-sga` filtering works: high heterozygosity allows the algorithm to perform well because the majority of read pairs carry at least one and often multiple sites informative about their haplotype phase. Therefore, the algorithm eliminates over 95% of heterozygosity in the read sets in the *Heliconius* trios, and the most striking improvement can be seen in the *Heliconius* cross. Still, it is also clear that the algorithm can substantially reduce heterozygosity even when the initial levels are moderate, as seen in the simulated yeast trio.

To assess the performance of the complete workflow we carried out full assemblies of all four trio datasets, as well as standard `sga` assemblies of the four offspring, and an assembly of one haploid *S. pombe* strain (JB759). Key statistics for all datasets are compared in Table 1. It is clear from the results that heterozygosity in the reads has substantial effect on the contiguity of assemblies, both at the contig and at the scaffold level. Applying `trio-sga` algorithms to the yeast ‘simulated trio’, where the offspring has heterozygosity of approximately 0.004 (1/262), improves the N50 measurement of contiguity by 69.3% at the contig level, and by 130% at the scaffold level. These are considerable improvements. To provide additional context for these numbers, the statistics for the yeast haploid assembly illustrate the further increases in contiguity that could be attained if all remaining heterozygous sites would be removed.

**Table 1:**
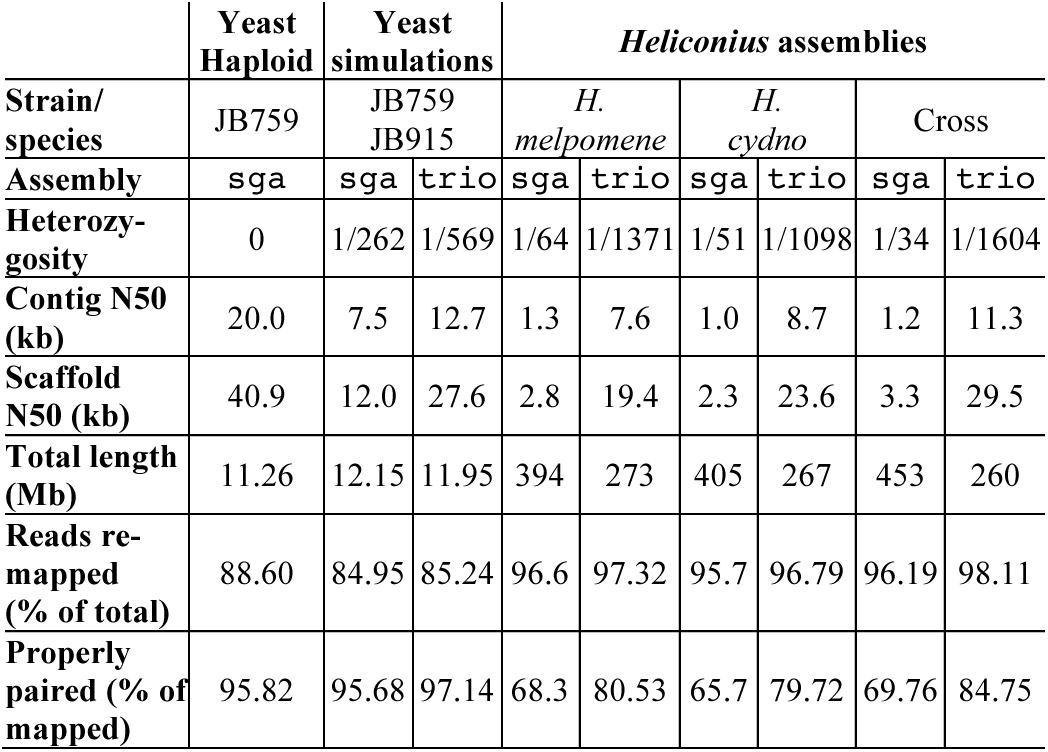
Assembly statistics for all datasets. Statistics for `trio-sga` refer to paternal haplotype assemblies. Contigs and scaffolds shorter than 500bp were excluded. All reads were re-mapped to the final scaffolds. Reads are properly paired if both map to the same scaffold, are correctly oriented, and their separation is within 5 standard deviations from the mean (i.e. the "proper pair" flag bit is set in the SAM file).

Normal assemblies of the highly heterozygous *Heliconius* genomes were very challenging for `sga`. Contig and scaffold N50 statistics are barely above 1kb and 2-3kb respectively. Moreover, the total length of these assemblies is much greater than the 292Mb flow cytometry genome size estimate for *H. melpomene* (Jiggins *et al.*, 2005), suggesting that in many cases two copies of a single genomic region have been retained. In contrast, trio assemblies have contig N50 between 7.5 and 11.2kb, scaffold N50 between 19.4 and 29.5kb, and total assembly lengths reflecting expected genome sizes. It is interesting to note that the most contiguous trio assemblies were for the *Heliconius* cross, again highlighting that `trio-sga` performance actually improves as heterozygosity in the initial dataset increases.

In all cases, the `trio-sga` assemblies have the higher proportion of reads-pairs that map back to the scaffolds as ‘properly paired’ (i.e. map to the same scaffold, are correctly oriented, and their separation is within 5 standard deviations from the mean), illustrating that the improvements in contiguity are not simply at the expense of assembly errors.

The distributions of scaffold lengths (Figure 5) provide a detailed depiction of contiguity of all the genome assemblies described in this manuscript. They reveal that the increases in N50 delivered by `trio-sga` are due to many improvements all across the scaffold length spectrum, rather than being due to a small number of long scaffolds.

**Fig. 5:**
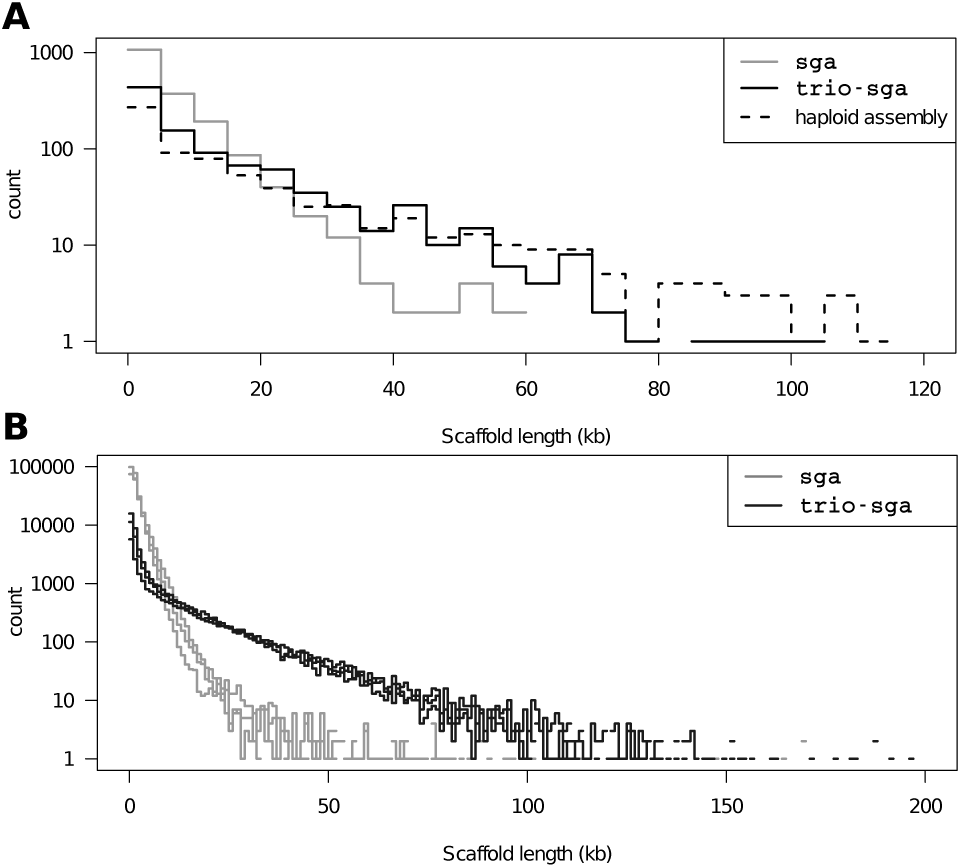
Scaffold length distributions. **(A)** The yeast assemblies, with scaffolds binned in 5kb intervals. **(B)** The six *Heliconius* assemblies (three `sga` and three with `trio-sga`), with scaffolds binned in 1kb intervals.

We note that further resolution of heterozygous sites and improvements in assembly contiguity may be achievable. The current `trio-sga` algorithms are able to use only a subset of the haplotype phase information present in the trio data. The additional cases, where the offspring and only one of the parents are heterozygous (loci 1 and 3 in Figure 2C) require identifying that the two variants are allelic. This may be identifiable in the assembly graph during the contig building process (i.e. at the stage corresponding to the `sga assemble` algorithm; Figure 1), but this is related to identifying whether branches correspond to repeats or heterozygous sites in the core assembly software, which is not the subject of this paper.

The ability of `sga` to scale to large genomes was demonstrated previously (Simpson and Durbin, 2012), and since has been substantially enhanced by further improvements in the speed and memory efficiency of FM-index construction algorithms (Bauer *et al.*, 2013); Heng Li’s implementation of the index construction has been used for the assemblies presented here (from github.com/lh3/ropebwt; available through `sga index-a ropebwt`).

The `trio-sga` subprogram that integrates the error-correction and phasing algorithms requires the forward FM-indices of all three members of the trio to be loaded into memory. Thus its memory requirement is approximately three times higher than required for standard sga read error-correction (an increase from 175MB to 503MB for yeast; and from 4.2GB to 12GB in *H. melpomene*). This step represents the highest point in terms of memory requirements. The CPU time required is also approximately three to four times higher (an increase from 3374 CPU sec. to 9441 CPU sec. in yeast; and from 55 CPU hours to 214 CPU hours in *H. melpomene*); however it can be run multithreaded on a single machine and further parallelized (for large gigabase scale genomes) by splitting the reads to be corrected and phased across multiple compute nodes. The subprogram for adding parents’ reads to the offspring assembly (`trio-sga filter-parents`) requires one third less memory and one third less CPU time.

## Acknowledgements

We thank Chris Jiggins and John Davey for supplying the *Heliconius* butterfly DNA, and specifically Richard Merrill and Ana Pinharanda for butterfly trio breeding and Sarah Barker for DNA extractions. We thank the Sanger Institute core facilities for sequencing. Raw reads from the Heliconius study are available from the European Nucleotide Archive (Study: ERP009507; Accessions: ERR926555-ERR926581). All the described genome assemblies are available from the Dryad Digital Repository (xxxxx-to be submitted on article acceptance).

## Funding

This work has been supported by the Wellcome Trust [097677/Z/11/Z to M.M., WT098051 to R.D.].

## Conflict of Interest

RD declares that he owns stock in Illumina from previous consulting.

## References

Adams, M.D. et al. (2000) The genome sequence of Drosophila melanogaster. Science, 287, 2185–2195.

Bauer, M.J. et al. (2013) Lightweight algorithms for constructing and inverting the BWT of string collections. Theoretical Computer Science.

Browning, B.L. and Browning, S.R. (2009) A unified approach to genotype imputation and haplotype-phase inference for large data sets of trios and unrelated individuals. Am. J. Hum. Genet., 84, 210–223.

C. elegans Sequencing Consortium (1998) Genome sequence of the nematode C. elegans: a platform for investigating biology. Science, 282, 2012–2018.

Charlesworth, D. and Willis, J.H. (2009) The genetics of inbreeding depression. Nat. Rev. Genet., 10, 783–796.

Donmez, N. and Brudno, M. (2011) Hapsembler: an assembler for highly polymorphic genomes. Research in Computational Molecular Biology.

Ferragina, P. and Manzini, G. (2000) Opportunistic data structures with applications. Foundations of Computer Science.

Genome 10K Community of Scientists (2009) Genome 10K: a proposal to obtain whole-genome sequence for 10,000 vertebrate species. J. Hered., 100, 659–674.

GIGA Community of Scientists et al. (2014) The Global Invertebrate Genomics Alliance (GIGA): developing community resources to study diverse invertebrate genomes. J. Hered., 105, 1–18.

i5K Consortium (2013) The i5K Initiative: advancing arthropod genomics for knowledge, human health, agriculture, and the environment. J. Hered., 104, 595–600.

Jeffares, D.C. et al. (2015) The genomic and phenotypic diversity of Schizosaccharomyces pombe. Nat. Genet., 47, 235–241.

Jiggins, C.D. et al. (2005) A genetic linkage map of the mimetic butterfly Heliconius melpomene. Genetics, 171, 557–570.

Kajitani, R. et al. (2014) Efficient de novo assembly of highly heterozygous genomes from whole-genome shotgun short reads. Genome Res., 24, 1384–1395.

Li, H. (2011) A statistical framework for SNP calling, mutation discovery, association mapping and population genetical parameter estimation from sequencing data. Bioinformatics, 27, 2987–2993.

Li, H. (2013) Aligning sequence reads, clone sequences and assembly contigs with BWA-MEM. arXiv, q-bio.GN.

Li, R.R. et al. (2010) De novo assembly of human genomes with massively parallel short read sequencing. Genes Dev., 20, 265–272.

Nystedt, B. et al. (2013) The Norway spruce genome sequence and conifer genome evolution. Nature, 497, 579–584.

O’Connell,J. et al. (2014) A general approach for haplotype phasing across the full spectrum of relatedness. PLoS Genet., 10, e1004234.

Pendleton,M. et al. (2015) Assembly and diploid architecture of an individual human genome via single-molecule technologies. Nat. Methods, 12, 780–786.

Potato Genome Sequencing Consortium et al. (2011) Genome sequence and analysis of the tuber crop potato. Nature, 475, 189–195.

Simpson,J.T. (2014) Exploring genome characteristics and sequence quality without a reference. Bioinformatics, 30, 1228–1235.

Simpson,J.T. and Durbin,R. (2010) Efficient construction of an assembly string graph using the FM-index. Bioinformatics, 26, i367–73.

Simpson,J.T. and Durbin,R. (2012) Efficient de novo assembly of large genomes using compressed data structures. Genome Res., 22, 549–556.

Vinson,J.P.J. et al. (2005) Assembly of polymorphic genomes: algorithms and application to Ciona savignyi. Genes Dev., 15, 1127–1135.

Zhang,G. et al. (2012) The oyster genome reveals stress adaptation and complexity of shell formation. Nature, 490, 49–54.

